# SARS-CoV-2 NSP12 associates with the TRiC complex and the P323L substitution is a host adaption

**DOI:** 10.1101/2023.03.18.533280

**Authors:** Muhannad Alruwaili, Stuart Armstrong, Tessa Prince, Maximilian Erdmann, David A. Matthews, Andrew Davidson, Waleed Aljabr, Julian A. Hiscox

## Abstract

SARS-CoV-2 emerged into the human population in late 2019 and human to human transmission has dominated the evolutionary landscape and driven the selection of different lineages. The first major change that resulted in increased transmission was the D614G substitution in the spike protein. This was accompanied by the P323L substitution in the viral RNA dependent RNA polymerase (RdRp) (NSP12). Together, with D614G these changes are the root of the predominant global SARS-CoV-2 landscape. Here, we found that NSP12 formed an interactome with cellular proteins. The functioning of NSP12 was dependent on the T-complex protein Ring Complex, a molecular chaperone. In contrast, there was differential association between NSP12 variants and components of a phosphatase complex (PP2/PP2A and STRN3). Virus expressing NSP12_L323_ was less sensitive to perturbations in PP2A and supports the paradigm that ongoing genotype to phenotype adaptation of SARS-CoV-2 in humans is not exclusively restricted to the spike protein.

## Introduction

The first major changes in the SARS-CoV-2 genome in the human population after spill overs from the intermediate animal host in the Huanan Seafood Wholesale Market (Pekar et al., 2022; Worobey et al., 2022) were the D614G substitution in the spike glycoprotein (spike_D614G_) accompanied by the P323L substitution in the viral RNA dependent RNA polymerase (NSP12) (NSP12_P323L_). These changes were associated with increased transmissibility and fitness (Plante et al., 2021) and mechanistically were linked to the spike_D614G_ substitution resulting in increased cell entry (Zhang et al., 2020). Whilst the spike_D614G_ substitution occurred in several different lineages, including lineage A, the combination of spike _D614G_ and NSP12_P323L_ substitutions were associated with the emergence of the B.1 lineage (Worobey et al., 2020) and descendent viruses that have come to dominant the global genomic landscape of SARS-CoV-2 (Andrews et al., 2022; Mlcochova et al., 2021; Volz et al., 2021). Focus on these changes was placed on spike_D614G_, with the NSP12_P323L_ substitution being considered a potentially fortuitous hitch hiker (Peacock et al., 2021). However, only the double mutation was epidemiologically successful and variants with single mutations, spike _D614_ a n d N S P_P3_1_23_2were a fraction of the global landscape (Ilmjarv et al., 2021). In support of NSP12 _P323L_ being biologically relevant, in animal models, viruses with NSP12 _L323_ had greater fitness than viruses with NSP12_P323_, suggesting a functional role in virus biology (Dong et al., 2021).

The role of the NSP12 _P323L_ substitution in SARS-CoV-2 biology has not been elucidated. In the virus cellular life cycle, NSP12 is the catalytic subunit of the polymerase complex. As such the protein does not function in isolation and forms a complex with (at least) two other viral proteins including the co-factors NSP7 and NSP8 (Kirchdoerfer and Ward, 2019; Wang et al., 2020). In a wider context the replication complex functions with other viral proteins that alter the structure of intracellular membranes (Wolff et al., 2020). The SARS-CoV-2 (and SARS-CoV) NSP12 is divided into several domains including the N-terminal nucleotidyl transferase (NiRAN) domain, the interface between amino acids 251 and 398, and the C-terminal polymerase domain. Structural analysis suggested that the interface could act as a protein interaction junction linking NiRAN with the fingers of the polymerase domain and NSP8 (Kirchdoerfer and Ward, 2019). The NSP12 _P323L_ substitution lays within the interface domain and is not obvious from structural analysis how this might alter the function of the protein.

We hypothesized that the NSP12_P323L_ substitution was a host adaptation that altered interaction with host cell proteins and functioned in virus RNA synthesis. Host proteins have been shown to interact with SARS-CoV-2 proteins (Gordon et al., 2020) and amino acid substitutions within these viral proteins may modulate their interaction with host proteins and function in the viral life cycle. Several RdRps from different viruses with RNA genomes interact with host proteins, including chaperones to promote stability and co-factors involved in replication (Camacho-Zarco et al., 2020; Munday et al., 2015; Taguwa et al., 2015).

To investigate this for SARS-CoV-2 NSP12, biochemical and virological approaches were combined to functionally characterise the interaction between NSP12_P323_ and NSP12_L323_ and the host cell proteome. The T-complex protein Ring Complex (TRiC) was identified as binding to both variants of NSP12 and disruption of chaperone activity ablated virus biology. In contrast, interactome analysis showed that protein phosphatase 2 (PP2/PP2A) associated more with NSP12 _L323_ rather than NSP12 _P323_. Reduction in the abundance of PP2A in virus infected cells resulted in a concomitant change in viral RNA synthesis but virus expressing NSP12_L323_ was less sensitive to these effects than virus expressing NSP12_P323_.

## Results

### Both the NSP12 _P323L_ variants interact with the TRiC complex but PP2A and STRN3 favour interaction with NSP12_L323_

To investigate the protein-protein interactions formed by NSP12 and whether there was a difference between NSP12_P323_ and NSP12 _L323_, an EGFP-based pull down approach was used in conjunction with mass spectrometry to identify and quantify potential cellular interactors followed by functional analysis for relevance to the virus life cycle (Figure 1A). Previously, we have used this approach to define the cellular interactome of a variety of different viral proteins, including the human respiratory syncytial virus RNA dependent RNA polymerase (L protein) (Munday *et al*., 2015), polymerase co-factors for Ebola virus (Dong et al., 2020) and the avian coronavirus nucleoprotein (Emmott et al., 2013). Many of these viral proteins retained biological activity with the EGFP tag. Here, either variant of NSP12 was fused to EGFP either C or N terminal and expressed in human cells in culture under the control of a CMV promoter (Figure 1A). The placing of the EGFP tag either N-terminal or C-terminal was to allow for any steric hindrance imparted by the EGFP moiety that might have affected protein-protein interactions. The plasmid constructs generated were pEGFP-NSP12_P323_, pNSP12_P323_-EGFP, pEGFP-NSP12 _L323_ and pNSP12 _L323_-EGFP. These plasmids were separately transfected into HEK 293T cells and expression confirmed by fluorescence, western blot and silver staining and compared to cells transfected with pEGFP only (expressing EGFP) (Figures 1B and C, respectively). This indicated that all four fusion proteins were expressed in cells, with C-terminal EGFP labelled proteins having a more punctate appearance (Figure 1B).

**Figure 1.**
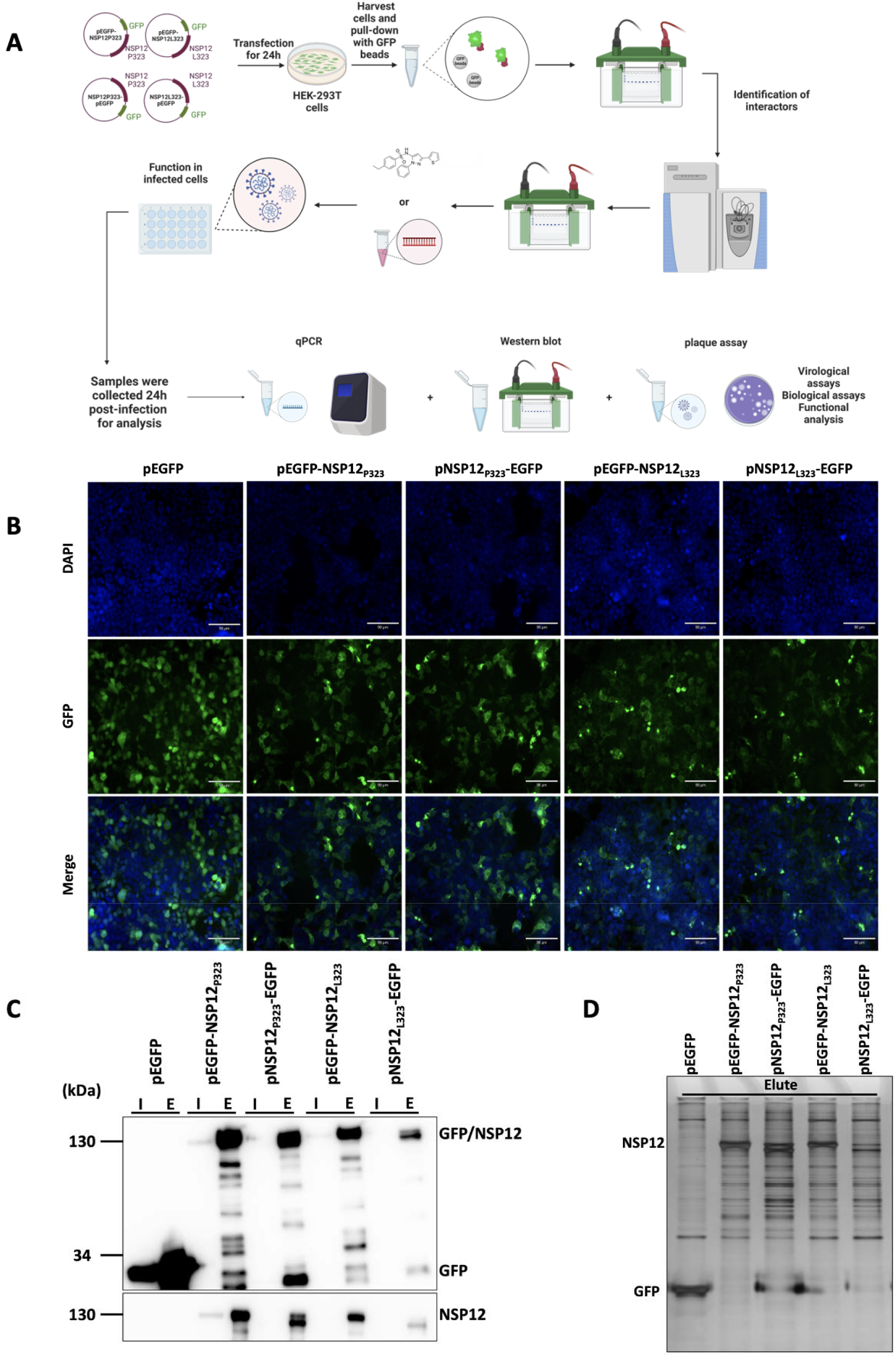
Expression of EGFP, EGFP-NSP12_P323_, NSP12 _P323_-EGFP, EGFP-NSP12 _L323_, and NSP12_L323_-EGFP in HEK-293T cells. . (A) Schematic diagram of the general methodology and approach used in this study. The NSP12 fusion proteins were expressed in HEK-293T cells and immunoprecipitated using a GFP-Trap. LC-MS/MS was used to identify the interactome and western blot used to validate key interactions. The functional implications of these findings were investigated using inhibitors or ablating the protein/function of interest and the effect on viral biology quantified. (B) Expression of EGFP, EGFP-NSP12 _P323_, NSP12 _P323_-EGFP, EGFP- NSP12_L323_, and NSP12 _L323_-EGFP in HEK-293T cells was confirmed by immunofluorescence microscopy with DAPI (blue) staining the nucleus. Scale bar, 90 μM. (C) Immunoblot analysis of input [I] and elute [E] EGFP, EGFP-NSP12_P323_, NSP12 _P323_-EGFP, EGFP-NSP12 _L323_, and NSP12_L323_-EGFP by western blot. Eluted samples for the mass spectrometry analysis were run on SDS-PAGE and stained with a silver stain, showing a difference in protein profile.

Western blot analysis using antibodies to either EGFP or NSP12 confirmed specific expression of the NSP12 moiety (Figure 1C), with overexposure suggesting the presence of breakdown/cleaved products. Silver stain analysis of protein lysates from pulldowns from the transfected cells highlighted the presence of the EGFP control and the NSP12 fusion proteins, with other protein species being present. These latter being suggestive of NSP12 interacting partners given the difference to EGFP only (Figure 1D).

To identify and quantify potential differences between the cellular interactomes of NSP12_P323_ or NSP12 _L323_, the five expression constructs were transfected five separate times into HEK 293T cells. Cells were lysed and the NSP12 moieties with EGFP and associated complexes immunoprecipitated using an EGFP-trap. Each pull down complex was characterised by mass spectrometry and cellular proteins compared across the different expression constructs. Criteria for inclusion as a positive ‘hit’ included identification of a protein by two or more unique peptides, scoring for each identified protein with a SAINT (Significance Analysis of INTeractome) (Choi et al., 2011) Bayesian FDR (BFDR) equal to or below 0.05. A MiST (Mass spectrometry interaction STatistics) (Verschueren et al., 2015) score of 0.6 and above was used as an additional filter to identify proteins unique to a particular construct. After this filtering process (BFDR ≤0.05), 91, 40, 987 and 50 proteins were identified as interacting with EGFP-NSP12_P323_, NSP12 _P323_-EGFP, EGFP-NSP12_L323_ and NSP12_L323_-EGFP, respectively (Supplementary Tables 1 to 4). The data indicated several common interactions across the four target proteins. These included components of the T-complex protein Ring Complex (TRiC) (or also known as Chaperonin Containing TCP-1 (CCT)), which is a multiple protein complex acting as a protein chaperone. However, some interactions were more enhanced/gained in EGFP-NSP12 _L323_ and NSP12_L323_-EGFP compared to EGFP-NSP12 _P323_ and NSP12 _P323_-EGFP. This included proteins serine/threonine-protein phosphatase 2A 65 kDa regulatory subunit A alpha isoform (PPP2R1A, a subunit of protein phosphatase 2 (PP2), also known as PP2A) and striatin-3 (STRN3) and epiplakin (EPPK1) (Figure 2). STRN3 can act as a regulatory subunit of PP2A (Tang et al., 2020). In cells, the function of PP2A is regulating phosphorylation and opposing the action of cellular kinases.

**Figure 2.**
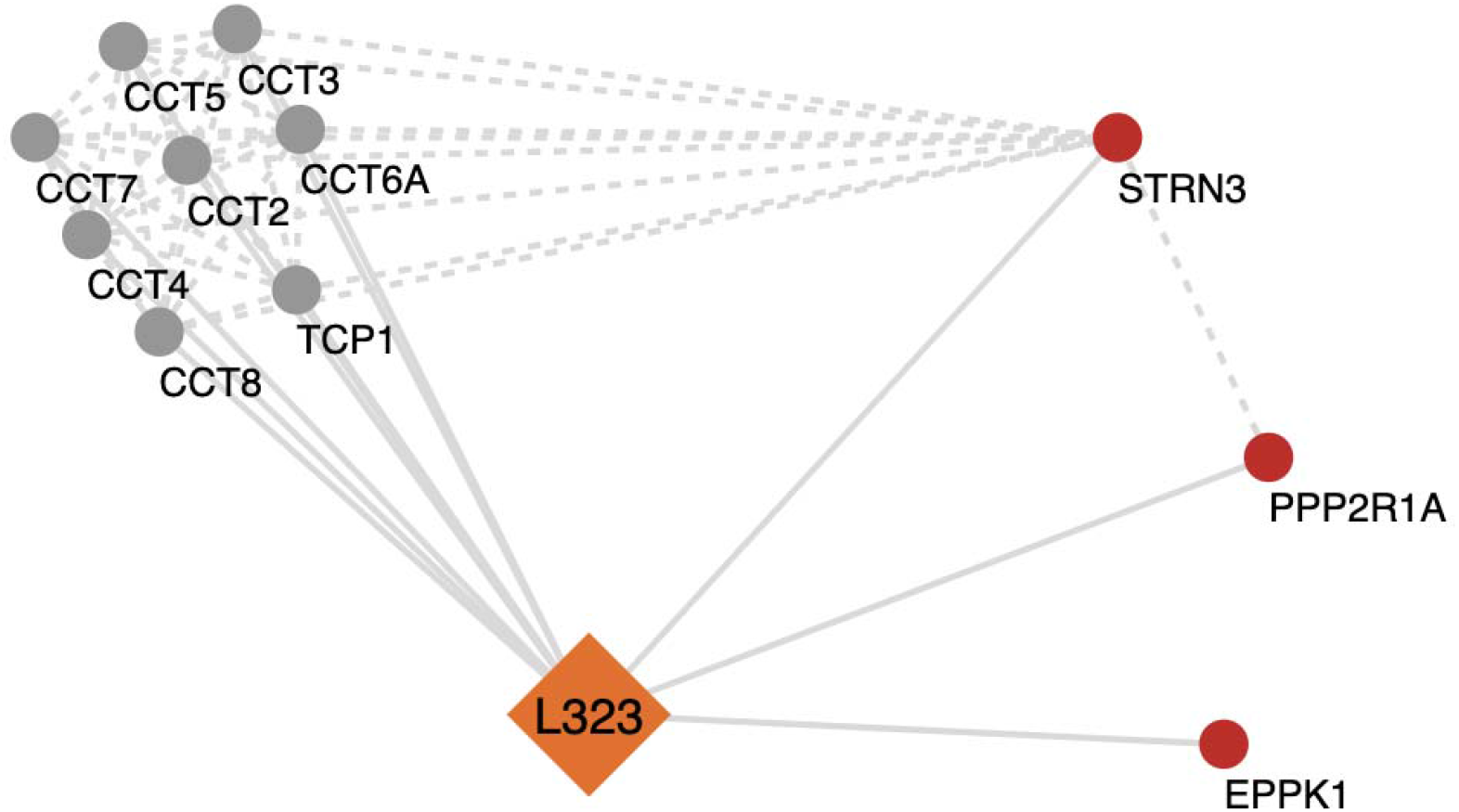
Analysis of selected significant cellular proteins that potentially interact with NSP12_L323_. Potential protein-protein interaction networks of NSP12 _L323_ were derived using Cytoscape from the mass spectrometry interactome data. Multiple proteins were found to be associated with NSP12_L323_. In this simplified figure, the highly enriched chaperonin containing TCP1 (TRiC/CCT complex) involved in the folding and assembly of proteins associated with NSP12_L323_ interactome. A few proteins were identified to be enhanced in the NSP12_L323_ interactome (STRN3 and PP2R1A which are part of the PP2A phosphatase family). Full interactome data for NSP12_P323_ and NSP12_L323_ are presented in Supplementary Figures 1 to 4 and Supplementary Tables 1 to 4.

### Validation of TRiC complex and PP2A and STRN3 interactions with NSP12_P323_ a n d NSP12_L323_

The interactome analysis indicated that NSP12 _L323_ interacted with the TRiC complex, which itself was linked to PP2A and STRN3 and these were subject to further investigation. Orthogonal techniques were used to validate the potential interaction between these cellular proteins and the viral targets. The transfection/expression experiments were repeated for each NSP12/EGFP fusion protein and using both forward (Figure 3A) and reverse pulldowns (Figure 3B) with specific antibodies to selected components of the TRiC complex - CCT1, CCT5, CCT7, CCT8 and also antibodies to PP2A and STRN3. The forward and reverse pulldowns confirmed the findings of the mass spectrometry analysis. Comparison of the cellular interactors of both variants of NSP12 showed that PP2A and STRN3 were more specific to EGFP tagged NSP12_L323_ than EGFP tagged NSP12_P323_. To investigate whether these interactions were mediated through binding to RNA (which may be plausible given NSP12 can associate with RNA), forward pulldowns were treated with RNase to remove species of cellular RNA (Figure 3C). Analysis of these treated pulldowns with the specific antibodies to the TRiC complex and PP2A and STRN3, indicated that none were mediated by RNA (Figure 3D).

**Figure 3.**
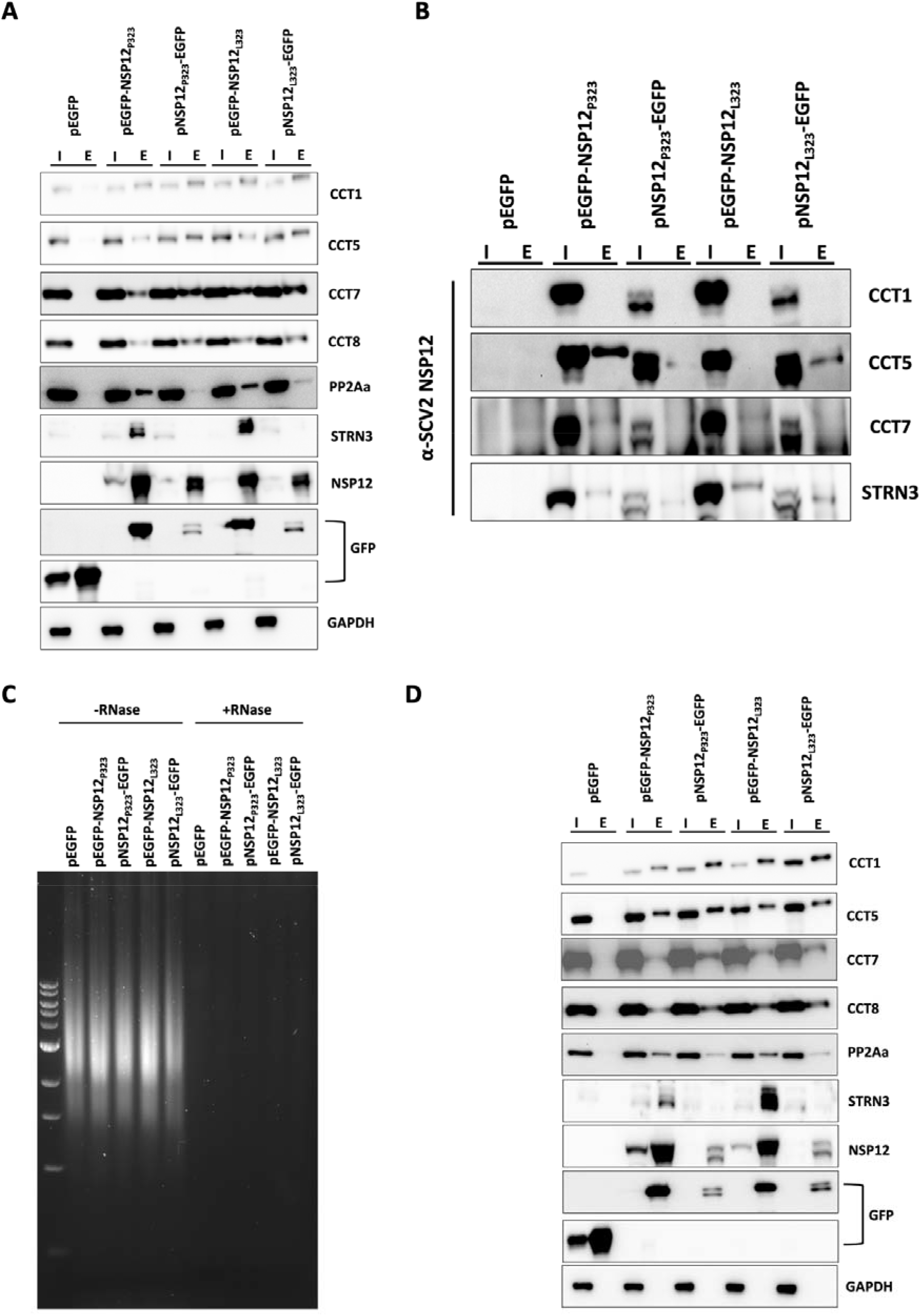
Validation of protein-protein interactions between EGFP tagged NSP12_P323_ or NSP12_L323_ and the host proteins in HEK 293T cells. (A) Transfection and pulldowns were repeated with EGFP, and EGFP tagged NSP12 _P323_, and NSP12 _L323_ at the C- and N- terminus for the detection of CCT1, CCT5, CCT7, CCT8, PP2Aa and STRN3 (I = input, E = elute). (B) Reverse pulldowns for CCT1, CCT5, CCT7, and STRN3 and detection of EGFP, EGFP-NSP12_P323_, NSP12_P323_-EGFP, EGFP-NSP12 _L323_, and NSP12 _L323_-EGFP with a specific antibody anti-NSP12. (C) Agarose gel electrophoresis showing that the treatment with RNase (+RNase) removed host RNA from cell lysate compared to untreated (-RNase). (D) Samples from (C) were run on SDS PAGE for the detection of host proteins in the absence of RNA using a forward pulldown approach.

### Functional role of the TRiC complex in the life cycle of SARS-CoV-2

To investigate whether the interaction between the TRiC complex and NSP12 was important for SARS-CoV-2 replication, the function of the TRiC complex was inhibited using the small molecule inhibitor HSF1A (heat shock transcription factor 1). A dose dependent assay was used to determine what concentration of the inhibitor could be used without affecting cell viability in either ACE2-A549 cells (used in infection assays – Supplementary Figure 5A) or HEK 293T cells (used in over expression analysis - Supplementary Figure 5B). The cell viability in the presence of different concentrations of HSF1A was similar between the two cell types, with 200 μM being equivalent to that induced by etoposide (which is a known agent/positive control that induces apoptosis). We hypothesized that disruption of the TRiC complex would have a negative impact on SARS-CoV-2 biology and disrupt NSP12.

Therefore, to test this hypothesis, the functioning of the TRiC complex was disrupted with 25 μM, 50 μM and 100 μM HSF1A in ACE2-A549 cells. These cells were infected with either recombinant SARS-CoV-2 expressing NSP12_P323_ or recombinant SARS-CoV-2 expressing NSP12_L323_. The effect of no inhibitor and inhibitor on virus biology was assessed. Several measures were used including RT-qPCR to determine the abundance of genomic and subgenomic RNA, the abundance of the nucleoprotein by western blot and comparison of viral titres. The data indicated that inhibition of the TRiC complex affected the biology of both SARS-CoV-2 expressing NSP12_P323_ (Figure 4A and B) or SARS-CoV-2 expressing NSP12_L323_ (Figure 4C and D). This resulted in a significant reduction in the expression of viral RNAs, viral protein and viral titres compared to virus grown in cells that were unexposed or received the vehicle only control (DMSO).

**Figure 4.**
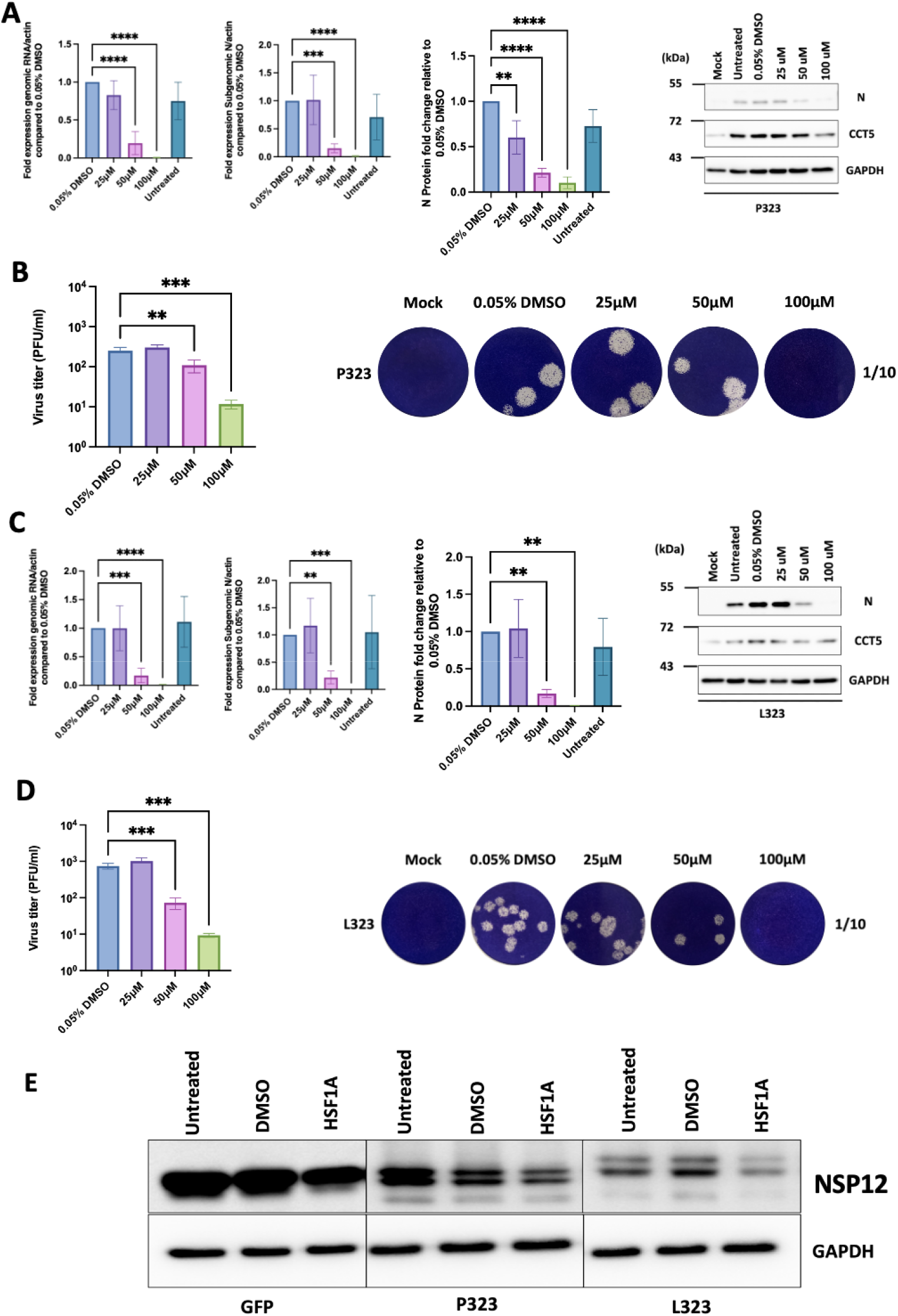
Determining the effect of the TRiC/CCT complex inhibitor HSF1A on the recombinant virus P323 and L323 replication. . (A and C) ACE2-A549 cells were pre-treated with either DMSO (control) or HSF1A (25, 50, or 100 µM) before being mock infected or infected with recombinant viruses expressing either NSP12_P323_ or NSP12 _L323_ viruses. At 24 hours after infection, gRNA and sgRNA levels were measured by qRT-PCR and normalised to β-actin. At the same time, whole cell lysate samples were collected, and the expression of nucleocapsid and CCT proteins was detected using western blot analysis with specific antibodies. GAPDH was used as a loading control. ImageJ was used to assess the expression of the nucleocapsid protein. Viral protein expressions were normalised to GAPDH. (B and D) ACE2-A549 cells were pre-treated for four hours with DMSO (control) or HSF1A (25, 50, and 100 µM) before being mock infected or infected with reverse genetically created P323 or L323 viruses at MOI of 0.1. Infected cells were maintained in media containing 0.05% DMSO or the previous concentrations of HSF1A for 24h. Viral titres were determined by plaque assay. (E) HEK-293T cells were transfected with a plasmid containing EGFP as a control, NSP12_P323_-EGFP, or NSP12 _L323_-EGFP respectively. Whole cell lysates were collected, and western blot analysis was performed to detect the abundance of EGFP, NSP12 _P323_-EGFP, and NSP12_L323_-EGFP proteins. GAPDH was used as a loading control. ** p<0.01, *** p<0.001, and **** p<0.0001 by one-way ANOVA with Dunnett’s multiple comparison test. All error bars represent the standard deviation.

The interaction between protein chaperones and viral encoded RdRps suggested that these complexes were involved in maintaining/facilitating the stability and function of the viral protein (e.g. (Munday *et al*., 2015; Taguwa *et al*., 2015)). Given the TRiC complex is involved in protein folding and stability, we hypothesised that inhibition of the TRiC complex would result in disruption of NSP12 and manifest in decreased abundance of the protein. To investigate this, NSP12 _P323_-EGFP or NSP12 _L323_-EGFP were overexpressed in HEK 293T cells exposed to HSF1A. The data indicated that in the presence of HSF1A, NSP12_P323_-EGFP and NSP12_L323_-EGFP were less abundant compared to expression of these proteins in cells exposed to vehicle only control (DMSO) or in the absence of vehicle/vehicle and inhibitor (Figure 4E). Taken together the data suggested that the inhibitor disrupted the activity of the TRiC complex and that this structure was important in maintaining the stability of NSP12.

### The NSP12_P323L_ substitution may act as a host adaptation mutation

The cellular interactomes, of EGFP-NSP12 _P323_ or EGFP-NSP12 _L323_, were identical in terms of interactions with cellular proteins (the TRiC complex being an example) but differed in degree of potential association with PP2Aa and STRN3. Mass spectrometry analysis indicated that these latter two proteins were more abundant in pulldowns where EGFP- NSP12_L323_ was used as a bait compared to EGFP-NSP12_P323_. To investigate whether these interactions had biological relevance, the mRNAs encoding STRN3 and PP2Aa were ablated using an siRNA-based approach (Supplementary Figure 6A and 6B, respectively), which resulted in a decrease in the abundance of both proteins after 48 hrs of exposure (Supplementary Figure 6C and 6D, respectively) with little impact on cell viability (Supplementary Figure 6E and 6F for STRN3 and PP2Aa, respectively). In both depletions a non-targeting siRNA and siRNA to GAPDH were used as a positive control to monitor knockdown and assess potential off target effects.

For the infection experiments, ACE2-A549 cells were depleted of STRN3 and PP2Aa for 48 hrs prior to infection with the recombinant viruses expressing either NSP12_P323_ or NSP12_L323_. Cells were infected for 24 hrs and viral biology assayed using RT-qPCR to detect and compare viral RNA, and western blot to identify the nucleoprotein, STRN3 and PP2Aa (Figure 5). The data indicated that depletion of STRN3 in cells infected with SARS-CoV-2 NSP12_P323_ resulted in no significant change in the abundance of viral RNAs, proteins or titre (Figure 5A and B). In contrast for cells infected with SARS-CoV-2 NSP12 _P323_ and depleted of PP2A, there was a significant reduction in the abundance of viral RNAs, proteins and titre (Figure 5A and B). In cells infected with SARS-CoV-2 NSP12_L323_ a slightly different pattern emerged. In cells depleted of STRN3 (Figure 5C), there was only a reduction in the abundance of the nucleocapsid protein sgRNA, the other virological factors were not significantly different. For the depletion of PP2A in cells infected with SARS-CoV-2 NSP12 _L323_ there was a significant reduction in the abundance of viral subgenomic mRNA (but the abundance of genomic RNA remained unchanged), nucleocapsid protein and viral titre. The major difference between SARS-CoV-2 NSP12 _P323_ and SARS-CoV-2 NSP12 _L323_ was that the latter variant was less sensitive to the depletion of PP2Aa than the former.

**Figure 5.**
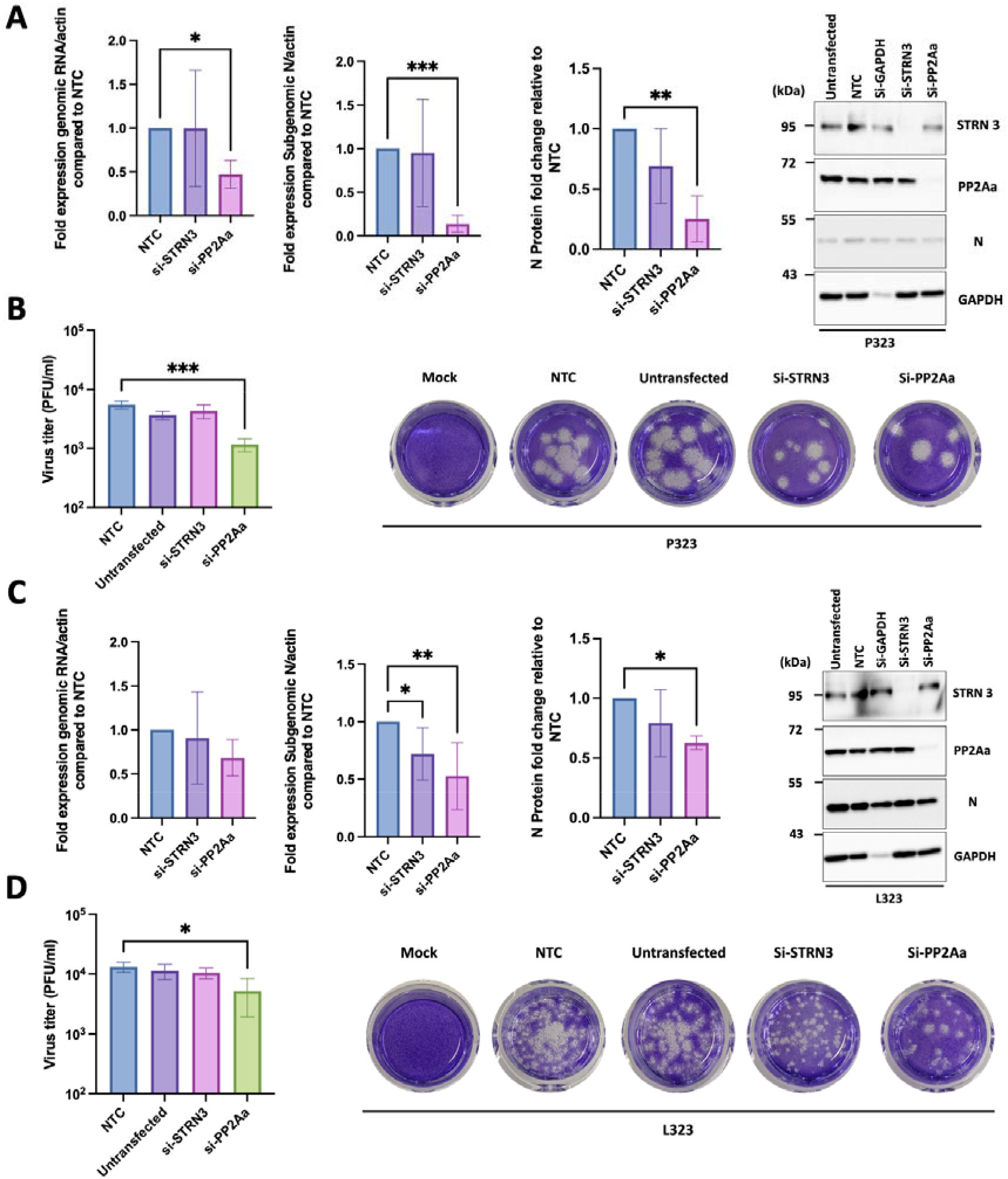
The impact of STRN3 and PP2Aa suppression on the replication of recombinant P323 and L323 viruses. (A and C) ACE2-A549 cells were mock transfected or transfected with non-targeting siRNA (NTC), si-GAPDH (to monitor transfection efficacy), si-STRN3, or si- PP2Aa for 48h. Subsequently, the ACE2-A549 cells were infected with recombinant viruses expressing either NSP12 _P323_ o r N S P_L3_1_23_2. RNA samples were extracted, and gRNA and sgRNA levels of the recombinant viruses were determined by qRT-PCR and normalised to β- actin. At the same time, whole cell lysate samples were collected, and the expression of nucleocapsid, STRN, and PP2Aa proteins was detected using western blot analysis. GAPDH was used as a loading control. ImageJ was used to assess the expression of the nucleocapsid protein. Viral protein expressions were normalised to GAPDH. (B and D) Supernatants, from the experiments described in panels A and C, were collected and viral titres were determined by plaque assay. * p<0.05, ** P<0.01, and *** p<0.001 by one-way ANOVA with Dunnett’s multiple comparison test. Data shown in (A-D) represent the mean of three independent experiments. (E) Data shown is from one experiment. All error bars represent the standard deviation.

## Discussion

SARS-CoV-2 emerged into the human population in late 2019 after several zoonotic jumps (Pekar *et a* ,*l .*2022; Worobey *et a l*, .2022) and presumably transiting through population bottlenecks. The first major genomic changes that resulted in different viral phenotypes was spike_D614G_ glycoprotein accompanied by NSP12 _P323L_. Several studies demonstrated that the spike_D614G_ substitution resulted in a gain in viral fitness including increased infectivity and stability of virions resulting in greater viral loads in the upper respiratory tract of people infected with SARS-CoV-2 (compared to contemporary viruses at the time) (Plante *et a* ,*l.* 2021). The NSP12 _P323L_ substitution was suggested to have occurred concurrently (Ilmjarv *et al*., 2021). Analysis of SARS-CoV-2 genomes in the containment and first lock down phase in the UK in 2020 showed rapid emergence of spike_D614G_ and NSP12 _P323L_ (Dong *et a l*,.2021). Infection experiments using in vivo models for COVID-19 suggested that the NSP12_P323L_ substitution also contributed a fitness advantage – over and above that provided by spike_D614G_ (Dong *et al*., 2021).

Whilst substitutions within SARS-CoV-2 may act at a structural level to stabilise RNA structure (Zhang et al., 2021) or stabilise viral proteins such as NSP12 (Periwal et al., 2022), these may also facilitate protein/protein interactions both between viral proteins but also with host proteins. Several studies have investigated the cellular interactome of the SARS-CoV-2 proteome including NSP12 (e.g. (Gordon *et a l*,.2020)). To investigate whether the P323L substitution was potentially host adapting we determined the cellular interactome of NSP12_P323_ and NSP12 _L323_ using an EGFP trap, by expressing N- and C-terminal fusion proteins. Whilst EGFP can be considered a larger moiety with the potential for steric hindrance we have found this a successful combination with a viral protein to assess expression and provide targeted enrichment for mass spectrometry, particularly in the absence of specific antibodies to the viral protein (Dong *et al*,. 2020; Munday *et al*,. 2015). The mass spectrometry analysis identified a potential cellular interactome of the two variants of NSP12. However, the high confidence hits identified in this study several in common with pre-printed studies (STRN3) (Bouhaddou et al., 2022) and components of the TRiC complex (but not TCP-1) (Laurent et al., 2020) but none in common to those previously reported for NSP12 in a published study (Gordon *et a* ,*l .*2020). This may be due to the differing bait/capture systems that were used and/or the filtering processes. In this study, selected interactions identified by the mass spectrometry analysis were validated with both forward and reverse pulldowns coupled to western blot (Figure 3A and B), but still with the presence of the EGFP tag. However, we were also able to demonstrate that the interactions were likely RNA independent (Figure 3D). Several components of the TRiC complex were identified as associating with NSP12. To our knowledge, whilst not described before for the coronavirus RdRp, interaction of TCP-1 with the influenza virus PB2 subunit of the RNA polymerase has been described and shown to be involved in replication (Fislova et al., 2010). The TRiC complex has also been shown to be involved in the replication of arenaviruses (negative sense segmented RNA viruses) (Sakabe et al., 2023). Pharmacological disruption of the TRiC complex in cells infected with SARS-CoV-2 expressing either NSP12_P323_ and NSP12 _L323_ (through reverse genetics) resulted in a 1-2 log fold reduction in viral titre depending on the variant (Figure 4). We note that in general, after 24hrs post-infection, the NSP12_L323_ variant tended to grow to a slightly higher titre in ACE2-A459 cells than the NSP12_P323_ variant, with concomitant slight differences in viral protein (nucleoprotein) and RNA abundances. Overall, we hypothesize that the TRiC complex stabilises the structure of NSP12 through chaperone activity, given that the abundance of an EGFP tagged NSP12 appeared to be lower in cells exposed to the TRiC inhibitor (Figure 4E). We cannot discount that this may be an off-target effect as the TRiC complex may also interact with other SARS-CoV-2 proteins critical to virus replication. However, we note that this has not been described in other interactome studies (Gordon *e t a*, *l*2.020). Ideally, we would have isolated the activity of NSP12 from other viral proteins. For example, in the context of TRiC complex disruption in a mini-replicon system. However, we have not had success recapitulating mini-replicon systems described in pre-printed studies.

Interestingly, two cellular proteins had a greater association with an EGFP-tagged N_L3_S_2_P_3_ 12 than the equivalent NSP12 _P323_ variant. These were STRN3, a regulatory subunit of the phosphatase, PP2Aa (Tang *et al*., 2020). We postulate that this is a host adapting mutation. Similar host adapting mutations/interactions have been described for the PB2 subunit of influenza virus (Camacho-Zarco *e t a*, *l*2.020). Likewise, the differential activity of the influenza virus polymerase varies between mammalian and avian cells contributing towards host range (Gabriel et al., 2007) and mutation of the polymerase complex led to adaptation in a new species (Gabriel et al., 2005; Taft et al., 2015).

Undoubtedly the D614G substitution in the spike protein resulted in a gain in fitness that manifested as a transmission advantage in humans. We would argue that the P323L substitution in NSP12 was not just a passive change and became dominant because it featured on genomes with G614 in spike, but itself conferred a fitness advantage and this may be due to enhanced interaction with cellular protein(s) involved in virus replication.

This would suggest that SARS-CoV-2 adaptation in humans is not just associated with changes in the spike protein and supports the paradigm that other viral proteins are involved (Thorne et al., 2022).

## Methods

### Cell Culture

Human Embryonic Kidney cells (HEK-293T) were maintained in DMEM with 10% FBS with no antibiotic and cells were used up to passage 30. Human ACE2-A549 (hACE2-A549), a lung epithelial cell line that overexpresses human ACE2 receptor, was kind gift of Oliver Schwartz (Buchrieser et al., 2020). Cells were maintained in DMEM with 10% FBS and 10ug/ml of Blasticidin (Invitrogen) to induce the overexpression of the ACE2 receptor. Passages 3 to 10 were used for experiments. All cell lines were tested regularly for mycoplasma contamination by PCR.

### Plasmid Design

SARS-CoV-2 (Wuhan sequence) was used for the construction of plasmids in this study. NSP12_P323_ and NSP12_L323_ sequences were codon-optimized for the expression in human cells. These sequences were synthesised at GeneArt and cloned into the pEGFP-C1 and pEGFP-N1 vectors. The resulting plasmids and inserts were confirmed by sequencing (data not shown).

### Plasmid Transfection

To obtain enough material for IP, 293T cells were seeded in two 145 cm^2^ dishes at 5x10^6^cells per dish 24h prior to calcium phosphate transfection. Cells were transfected with 25.6 μg of plasmid DNA encoding pEGFP, pEGFP-NSP12 _P323_, pNSP12 _P323_-EGFP, pEGFP-NSP12 _L323,_ and pNSP12 _L323_-EGFP. The cells were harvested 24hrs post-transfection, lysed, and immunoprecipitated using a GFP-Trap (Chromotek).

### EGFP Co-immunoprecipitations

pEGFP, pEGFP-NSP12_P323_, pNSP12_P323_-EGFP, pEGFP-NSP12_L323_, and pNSP12_L323_-EGFP transfected cells were harvested using a cell scraper and transferred to a 50 ml tube. Cells were pelleted at 1000 x g for 5 min followed by three washes with 10 ml of PBS. Cell pellets were resuspended in 200 μl of lysis buffer (10 mM Tris/Cl pH 7.5; 150 mM NaCl; 0.5 mM EDTA; 0.5% NP40) supplemented with Halt Protease Inhibitor Cocktail EDTA-Free (Thermo Scientific) and incubated for 30 min on ice. Lysates were transferred to 1.5 ml tubes and clarified by centrifugation at 14,000 x g, and the supernatants were transferred to new tubes and diluted five-fold with dilution buffer. GFP-Traps were equilibrated with ice-cold dilution buffer, and diluted cell lysate incubated with equilibrated beads on a rotator overnight at 4°C. Beads containing proteins were washed and centrifuged at 2,500 x g three times for two minutes to remove non-bound proteins. Proteins were eluted from beads through heating in 1x sample buffer at 85°C for 10 minutes. The solution was centrifuged at 2, 500 x g for two minutes, and the supernatants transferred to protein low bind tubes and stored at -80°C for further analysis.

### Reverse Co-immunoprecipitation

The immunoprecipitations using antibodies against CCT1, CCT5, CCT7, and STRN3 were carried out utilising immobilised recombinant protein A/G resin (Thermo Scientific). Cell pellets were harvested and processed as described in the EGFP co-immunoprecipitation section. Cell lysates were incubated with a concentration of antibodies recommended by the supplier for 2hrs on a rotator at 4°C. Protein A/G resin (50 μl) was equilibrated with ice-cold dilution buffer, incubated overnight at 4°C on a rotator with diluted cell lysate containing the antibody, and then centrifuged at 2,500 x g for two minutes to remove non-bound proteins. Then, the A/G protein resin complexed with the target protein was washed and eluted as previously described in EGFP co-immunoprecipitation.

### Protein digestion

In-gel digestion was performed as described (Shevchenko et al., 2006). Eluted proteins (20µl, in reducing sample buffer) were run approx. 1 cm into a 4-12% NuPage gel (Invitrogen) before staining with Coomassie blue (GelCode Blue Safe Protein Stain, Fisher) for at least 1 hr, then de-stained with ultrapure water for at least 2 hrs. The entire lane length (1mm wide) was excised and cut into smaller pieces (approx. 1mm^3^) before de- staining with 25mM ammonium bicarbonate/ 50 % acetonitrile (v/v). Proteins were reduced for 10 mins at 60°C with 10 mM dithiothreitol (Sigma) in 25 mM ammonium bicarbonate and then alkylated with 55 mM iodoacetamide (Sigma) in 50 mM ammonium bicarbonate for 30 mins in the dark at room temperature. Gel pieces were washed for 15 mins in 50 mM ammonium bicarbonate and then dehydrated with 100% acetonitrile. Acetonitrile was removed and the gel plugs rehydrated with 0.01 µg/µL proteomic grade trypsin (Thermo) in 25 mM ammonium bicarbonate. Digestion was performed overnight at 37°C. Peptides were extracted with 50% (v/v) acetonitrile, 0.1% TFA (v/v) and the extracts were reduced to dryness using a centrifugal vacuum concentrator (Eppendorf) and re-suspended in 3 % (v/v) methanol, 0.1 % (v/v) TFA for analysis by MS.

### NanoLC MS ESI MS/MS analysis

LC-MS/MS analysis was performed similar to that described by Aljabr *et al* (2019). Peptides were analysed by on-line nanoflow LC using the Ultimate 3000 nano system (Dionex/Thermo Fisher Scientific). Samples were loaded onto a trap column (Acclaim PepMap 100, 2 cm × 75 μm inner diameter, C18, 3 μm, 100 Å) at 9μl /min with an aqueous solution containing 0.1 %(v/v) TFA and 2% (v/v) acetonitrile. After 3 mins, the trap column was set in-line an analytical column (Easy-Spray PepMap® RSLC 50 cm × 75 μm inner diameter, C18, 2 μm, 100 Å) fused to a silica nano-electrospray emitter (Dionex). The column was operated at a constant temperature of 35°C and the LC system coupled to a Q-Exactive mass spectrometer (Thermo Fisher Scientific). Chromatography was performed with a buffer system consisting of 0.1 % formic acid (buffer A) and 80 % acetonitrile in 0.1 % formic acid (buffer B). The peptides were separated by a linear gradient of 3.8 – 50 % buffer B over 30 minutes at a flow rate of 300 nl/min. The Q-Exactive was operated in data-dependent mode with survey scans acquired at a resolution of 70,000 at m/z 200. Scan range was 300 to 2000m/z. Up to the top 10 most abundant isotope patterns with charge states +2 to +5 from the survey scan were selected with an isolation window of 2.0Th and fragmented by higher energy collisional dissociation with normalized collision energies of 30. The maximum ion injection times for the survey scan and the MS/MS scans were 250 and 50 ms, respectively, and the ion target value was set to 1x10^6^ for survey scans and 1x10^4^ for the MS/MS scans. MS/MS events were acquired at a resolution of 17,500. Repetitive sequencing of peptides was minimized through dynamic exclusion of the sequenced peptides for 20 secs.

### Protein Identification

MS spectra data was analysed by label-free quantification using MaxQuant software (version 1.6.17.0, (Cox et al., 2014)) and searched against the human protein database (Uniprot UP000005640_9606, May2021) and the SARS-CoV-2 NSP12 bait protein (NCBI Reference Sequence: YP_009725307.1). Detected proteins were filtered to 1% false discovery rate. MaxQuant results were further processed with SAINTexpress (version 3.6.3, Teo *et al* 2014) using default settings. High confidence interactions were classified as having a SAINTexpress Bayesian false-discovery rate (BFDR) ≤0.05, and an average spectral count ≥2. Additional filtering to highlight unique interactors was performed using MiST in HIV trained mode (Verschueren et al., 2015). Proteins with scores above 0.6 were deemed unique to that bait. PPI networks were generated in Cytoscape (Shannon et al., 2003) with protein-protein interactions and functional enrichment performed using STRING (Doncheva et al., 2019).

### Cell Viability Assay

Cell viability was measured using the CellTiter-Glo Luminescent Cell Viability Assay (Promega). Cells were seeded in opaque-walled clear 96 well plates (Corning) in triplicate for each condition. The plate was equilibrated at room temperature for 30 minute and 100 μl of the reagent was added. The plate was shaken at 450 rpm for two mins and incubated for 10 mins at room temperature. Luminescence was measured using the GloMax Explorer Reader (Promega) with an integration time of 0.3 secs per well. All values were normalised to the control, which was set to 100%.

### siRNA Knockdown

Lipofectamine (Invitrogen) (1.4 μl of RNAiMAX) was added to 23.6 μl of Opti-MEM I reduced serum media in a 1.5 ml tube. In a separate tube, the final concentration of a non-targeting siRNA or targeting siRNA was used at a concentration of 10 nM per well in a 25 μl volume of Opti-MEM I reduced serum media. siRNA was added onto the RNAiMAX and incubated for 5 mins at room temperature. The incubated complex (50 μl) was added to each well and hACE2-A549 cells in complete media in the absence of antibiotic were seeded at 0.05 x 10 ^6^ and incubated for 48 hrs at 37^°^C and in 5% CO _2_. Whole cell lysates were collected and analysed by western blot analysis to measure the knockdown efficiency.

### Virus Infection

Cells of ACE2-A549 were inoculated with either recombinant viruses expressing NSP12_P323_ or NSP12_L323_ for 1 hr at ^°^3C7and in 5% CO _2_. The inoculum was removed and replaced with DMEM containing 2% FBS with no antibiotic, or DMEM containing a different dose of HSF1A for TRiC inhibition study, for 24 hrs.

### Quantitative Real-Time PCR

RNA from cells infected with the virus was extracted with TRIzol reagent (Fisher) according to the manufacture’s protocol. Extracted RNA was then subjected to Turbo DNase treatment (Invitrogen). RNA (200 ng) was reverse transcribed into cDNA using LunaScript®RT SuperMix (NEB) according to the manufacturer’s protocol. Generated cDNA was diluted 1:4 in Nuclease free water. Primers (White et al., 2021) targeting SARS-CoV-2 gRNA and N protein sgRNA were used for the quantification by RT-qPCR using iTaq Universal SYBER Green Supermix (Bio-Rad) and normalised to β-actin using 2^-ΔΔCT^.

### Western Blot Analysis

Cells were lysed in either RIPA buffer or IP lysis buffer (for ACE2-A549 infected cells or HEK- 293T transfected cells) in the presence of a protease inhibitor cocktail. For ACE2-A549, 5 μg of whole cell lysate was heated at 70°C in 4X sample buffer for 10 minutes then separated by 10% SDS-PAGE for 1 hr. Proteins were transferred onto PVDF membrane. 5% non-fat milk in 0.1% TBST was used to block the membrane for 1 hr at room temperature. The blots were then incubated with primary antibodies overnight at 4°C and secondary antibodies for 1 hr at room temperature. Blots were washed in 0.1% TBST 5 mins three times each between each antibody. The blots were imaged using a chemiluminescent reagent (ECL) with ChemiDoc gel imaging system (Bio-Rad).

### Plaque Assay

A plaque assay was used to determine the virus titre of the supernatant collected from each condition. 10-fold dilutions of each supernatant were used to inoculate ≈ 90% confluent Vero E6 cells in duplicate for 1 hr at 37^°^C and 5% CO _2_. The inoculum was removed and replaced with DMEM containing 2% FBS and 2% low-melting-point agarose and incubated at 37^°^C and 5% CO _2_ for 72 hrs. Formalin (10%) was added to each well for 1 hr. Overlay containing formalin was removed and plates stained with crystal violet.

### Statistical Analysis

All the analysis (except the mass spectrometry data) were performed using GraphPad Prism software. One-way ANOVA test using Dunnett’s multiple comparison test. The significance level in this study was set at p-value of less than 0.05. All error bars represent the standard deviation.

## Acknowledgements

This work was funded by U.S. Food and Drug Administration Medical Countermeasures Initiative contract (75F40120C00085)1to JAH with Co-Is, ADD, WA and DAM. The article reflects the views of the authors and does not represent the views or policies of the FDA.1 This work was supported by the MRC (MR/W005611/1) G2P-UK: A national virology consortium to address phenotypic consequences of SARS-CoV-2 genomic variation (co-Is ADD and JAH). The work was supported by Northern Border University, Saudi Arabia.

## Author contributions

Conceptualization: MA and JAH. Methodology: MA, SA, TP, ME. Validation: MA, SA, TP, ME. Formal analysis: MA, ME and SA. Investigation: MA, SA, TP and ME. Resources: SA, ME, DAM and AD. Data curation: SA. Writing - Original Draft: MA and JAH. Writing - Review & Editing: all authors. Visualization: MA and SA. Supervision: DAM, AD, WA and JAH. Project administration: JAH. Funding acquisition: DAM, AD, WA and JAH.

---

**Supplementary Figure 1.**
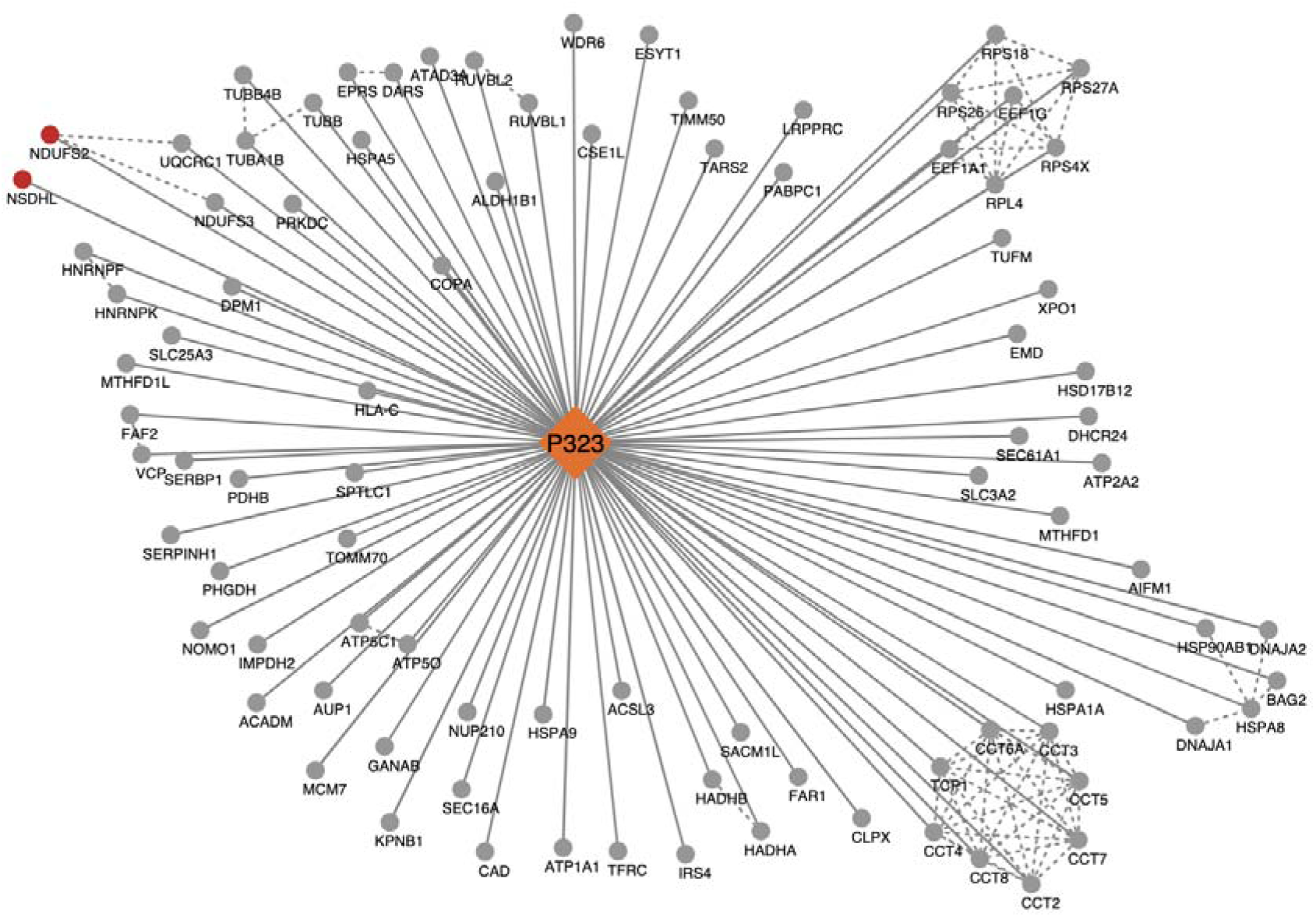
Host Protein Interactome with EGFP-NSP12_P323_. The putative protein-protein interaction network of NSP12 _P323_ with an n-terminal GFP tag was obtained via affinity pulldown mass spectrometry (AP-MS). PPI networks were created using cytoscape and STRING. The grey nodes denote high confidence interacting proteins (SAINTexpress BFDR score ≤ 0.05). The red nodes denote proteins that are unique to this NSP12 construct (BFDR score ≤0.05, MiST score ≥0.6). Solid lines represent putative interaction with the bait protein. Dashed lines represent previously known associations obtained via STRING. Proteins with enhanced association with EGFP-NSP12_P323_ include NSDHL (Sterol-4-alpha-carboxylate 3-dehydrogenase) which is involved in sterol synthesis and NDUFS2 (NADH dehydrogenase [ubiquinone] iron-sulfur protein 2), a mitochondrial protein subunit in Respiratory complex I. Protein clusters involved with translation (RPS27A, RPL4, EEF1G, EEF1A1, RPS7, RPL3, RPS26, RPS4X, RPL11, RPS18, RPL10, RPS19) and protein folding (TRiC complex,CCT-CCT8, TCP1) are highly enriched.

**Supplementary Figure 2.**
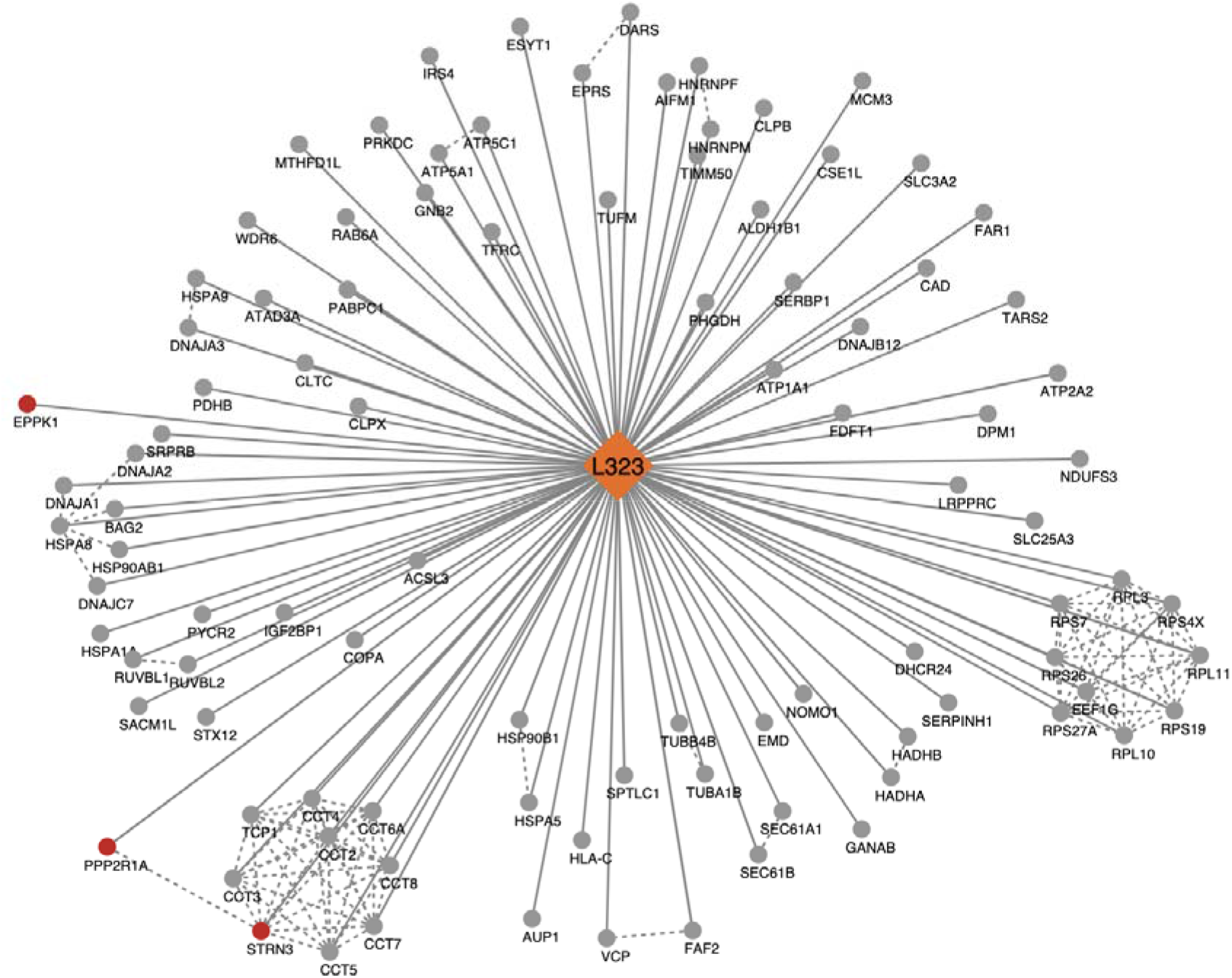
Host Protein Interactome with EGFP-NSP12_L323_. The putative protein-protein interaction network of NSP12 _L323_ with an n-terminal GFP tag was obtained via affinity pulldown mass spectrometry (AP-MS). PPI networks were created using cytoscape and STRING. The grey nodes denote high confidence interacting proteins (SAINTexpress BFDR score ≤ 0.05). The red nodes denote proteins that are unique to this NSP12 construct (BFDR score ≤0.05, MiST score ≥0.6). Solid lines represent putative interaction with the bait protein. Dashed lines represent previously known associations obtained via STRING. Proteins with enhanced association with EGFP-NSP12_L323_ include EPPK1 (Epiplakin), which has roles in reorganization of the cytoskeleton and cell proliferation and STRN3 (striatin-3) and PPP2R1A (Serine/threonine-protein phosphatase 2A) which are part of the PP2A phosphatase family. Protein clusters involved with translation (RPS27A, RPL4, EEF1G, EEF1A1, RPS7, RPL3, RPS26, RPS4X, RPL11, RPS18, RPL10, RPS19) and protein folding (TRiC complex, CCT-CCT8, TCP1) are highly enriched.

**Supplementary Figure 3.**
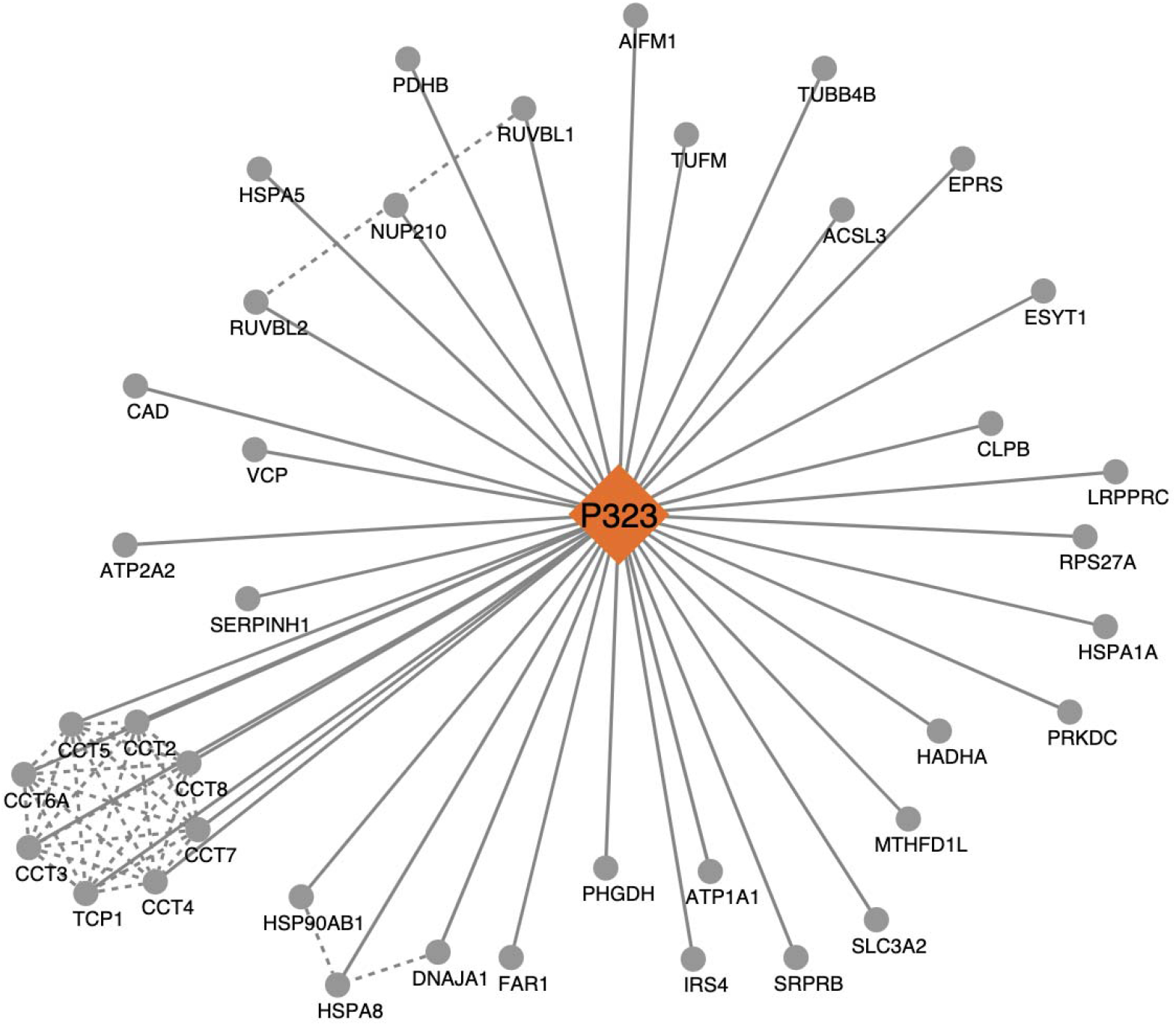
Host Protein Interactome with NSP12_P323_-EGFP. The putative protein-protein interaction network of NSP12 _P323_ with an c-terminal GFP tag was obtained via affinity pulldown mass spectrometry (AP-MS). PPI networks were created using cytoscape and STRING. The grey nodes denote high confidence interacting proteins (SAINTexpress BFDR score ≤ 0.05). The red nodes denote proteins that are unique to this NSP12 construct (BFDR score ≤0.05, MiST score ≥0.6). Solid lines represent putative interaction with the bait protein. Dashed lines represent previously known associations obtained via STRING. There are no proteins with enhanced association with NSP12_P323_. Protein cluster involved with protein folding (TRiC complex, CCT-CCT8, TCP1) is highly enriched.

**Supplementary Figure 4.**
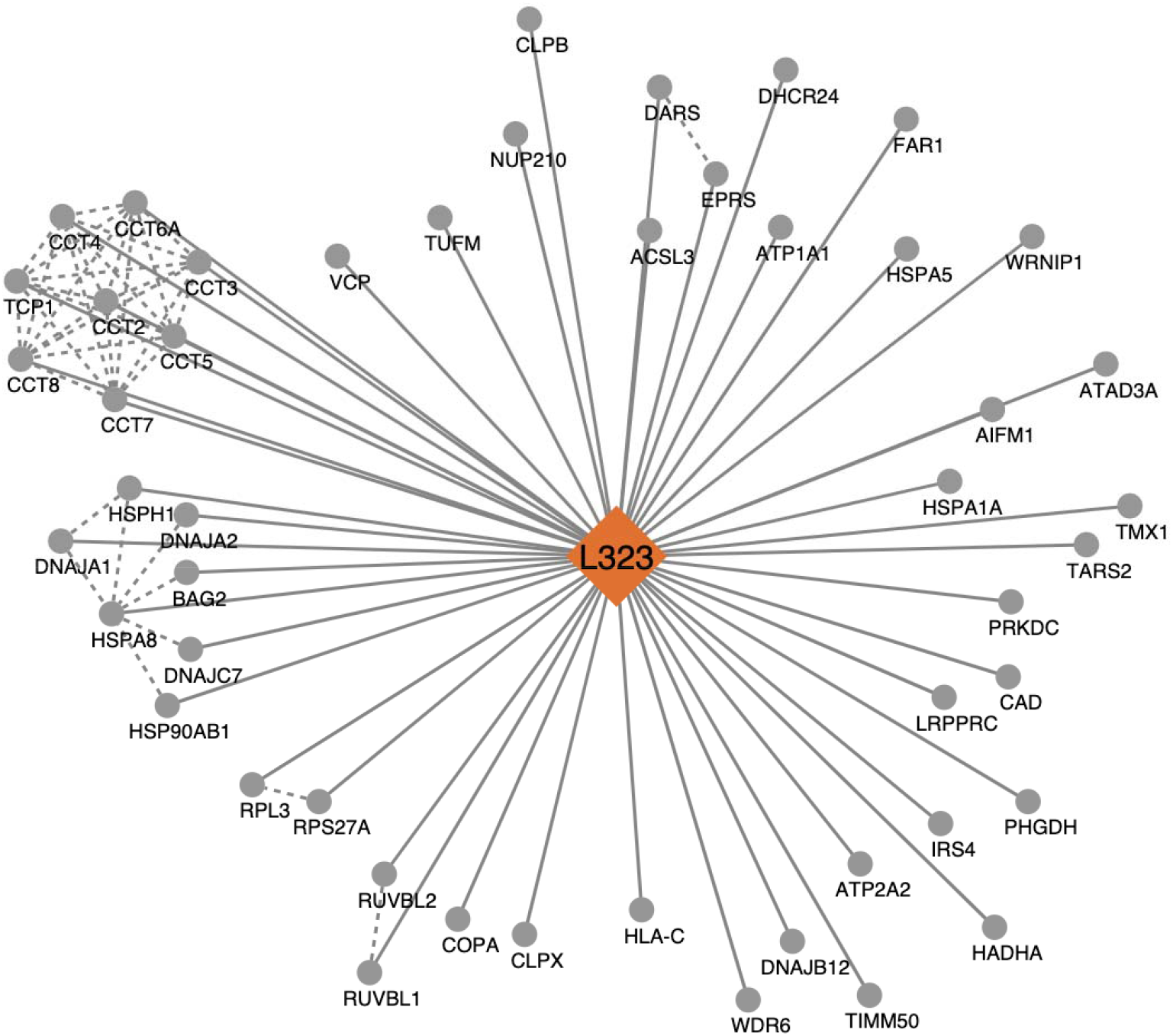
Host Protein Interactome with NSP12_L323_-EGFP. The putative protein-protein interaction network of NSP12 _L323_ with an c-terminal GFP tag was obtained via affinity pulldown mass spectrometry (AP-MS). PPI networks were created using cytoscape and STRING. The grey nodes denote high confidence interacting proteins (SAINTexpress BFDR score ≤ 0.05). The red nodes denote proteins that are unique to this NSP12 construct (BFDR score ≤0.05, MiST score ≥0.6). Solid lines represent putative interaction with the bait protein. Dashed lines represent previously known associations obtained via STRING. There are no proteins with enhanced association with NSP12_L323_. Protein cluster involved with protein folding (TRiC complex, CCT-CCT8, TCP1) is highly enriched.

**Supplementary Figure 5.**
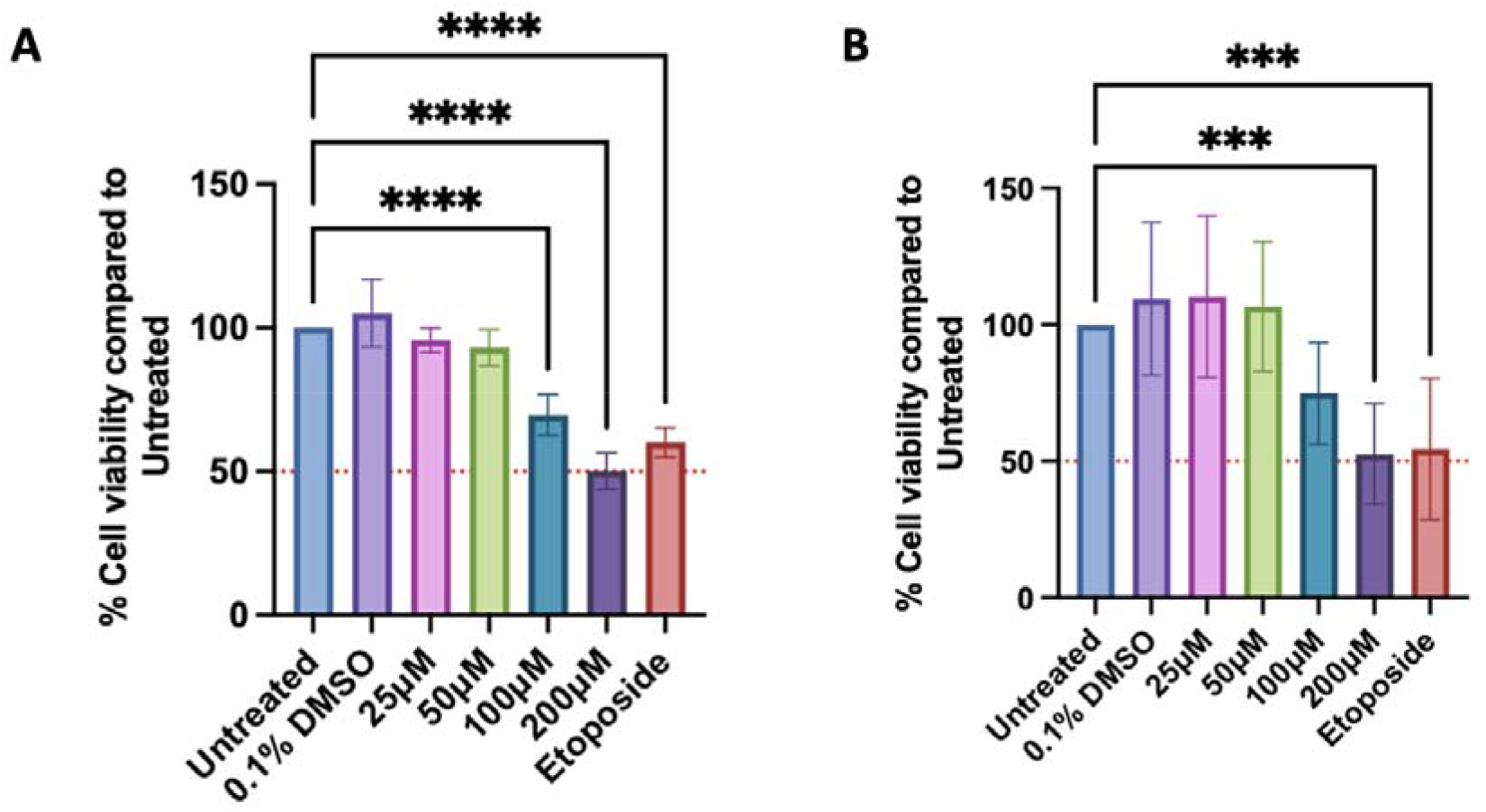
Cell viability assay (CellTiter Glo assay) of ACE2-A549 and HEK- 293T cells treated with the TRiC/CCT complex inhibitor, HSF1A. . (A) ACE2-A549 or (B) HEK- 293T cells were treated with a range of HSF1A concentrations from 25 to 200 μM for 28 h. Untreated cells were used as a control; 60 µM of Etoposide was used as a positive control for the induction of cell death. The luminescence for untreated cells was normalised to 100%, then % cell viability for different HSF1A doses were calculated relative to this. The dashed red line indicates the GI50. *** p<0.001 and **** p<0.0001 by one-way ANOVA with Dunnett’s multiple comparison test. Data shown represent the mean of three independent experiments. All error bars represent the standard deviation.

**Supplementary Figure 6.**
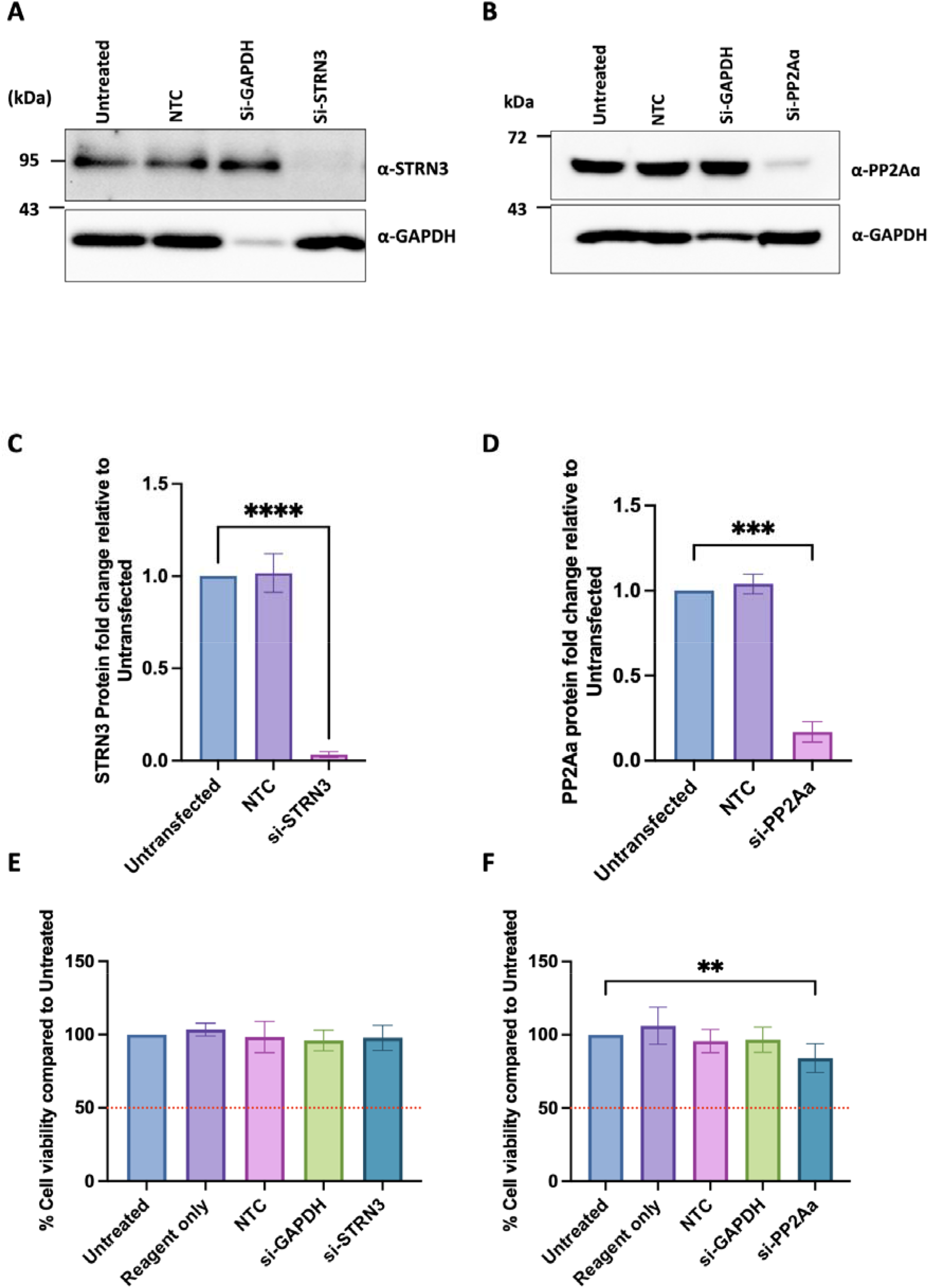
STRN3 and PP2Aa siRNA knockdown efficiency. (A) ACE2-A549 cells were reverse-transfected with 10 nM of either non-targeting siRNA control (NTC) alongside si-GAPDH (to monitor knockdown efficiency) and si-STRN3. After 48 h, whole cell lysates were collected and evaluated for the expression levels of STRN3 by western blot analysis. (B) ACE2-A549 cells were reverse-transfected with 10 nM of either non-targeting siRNA control (NTC) alongside si-GAPDH (to monitor knockdown efficiency) and si-PP2Aa. After 48 h, whole cell lysates were collected and evaluated for the expression levels of PP2Aa by western blot analysis. ImageJ quantification of (C) STRN3 or (B) PP2Aa expressions and normalised to the internal control GAPDH. ACE2-A549 cells were reverse-transfected in 96 well plate format with 10 nM of either non-targeting siRNA control (NTC) alongside si- GAPDH (E) si-STRN3, or (F) si-PP2Aa for 48 h. the mean luminescence for untreated cells was normalised to 100%, and then % cell viability for different conditions were calculated relative to this. Data shown represent the mean of three independent experiments. All error bars represent the standard deviation.

